# Analysis of the human kinome and phosphatome reveals diseased signaling networks induced by overexpression

**DOI:** 10.1101/314716

**Authors:** Xiao-Kang Lun, Damian Szklarczyk, Attila Gábor, Nadine Dobberstein, Vito RT Zanotelli, Julio Saez-Rodriguez, Christian von Mering, Bernd Bodenmiller

## Abstract

Kinase and phosphatase overexpression drives tumorigenesis and drug resistance in many cancer types. Signaling networks reprogrammed by protein overexpression remain largely uncharacterized, hindering discovery of paths to therapeutic intervention. We previously developed a single cell proteomics approach based on mass cytometry that enables quantitative assessment of overexpression effects on the signaling network. Here we applied this approach in a human kinome- and phosphatome-wide study to assess how 649 individually overexpressed proteins modulate the cancer-related signaling network in HEK293T cells. Based on these data we expanded the functional classification of human kinases and phosphatases and detected 208 novel signaling relationships. In the signaling dynamics analysis, we showed that increased ERK-specific phosphatases sustained proliferative signaling, and using a novel combinatorial overexpression approach, we confirmed this phosphatase-driven mechanism of cancer progression. Finally, we identified 54 proteins that caused ligand-independent ERK activation with potential as biomarkers for drug resistance in cells carrying BRAF mutations.

## Introduction

Kinases and phosphatases control the reversible process of phosphorylation, which regulates protein structure, activity, and localization. Kinases and phosphatases are organized as signaling networks that compute extracellular signals into transcriptional, functional, and phenotypical responses. Deregulation of signaling networks can lead to the initiation and progression of many types of human disease including cancer (Fleuren et al., 2016; Julien et al., 2007), consequently they are a focal point of life science research. Kinases and phosphatases have been classified based on genomic and proteomic analyses (Chen et al., 2017; Manning et al., 2002; Sacco et al., 2012a). Signaling network structure has been studied by mapping physical interactions of kinases and phosphatases in steady and dynamic states using biochemical approaches and reporter assays in yeast and human cells (Barrios-Rodiles et al., 2005; Breitkreutz et al., 2010; Couzens et al., 2013; Horn et al., 2011). Using *in vitro* kinase assays and motif-based predictions, the specificity and targets of many kinases have been revealed (Linding et al., 2007; Mok et al., 2010; Yu et al., 2009). Kinase and phosphatase perturbations have also been applied to systematically determine phosphorylation abundance changes in yeast and human cells (Bodenmiller et al., 2010; Ochoa et al., 2016; Sacco et al., 2012b). A systematic analysis of how each human kinase and phosphatase modulates signaling network structure and dynamics, however, so far is absent.

Mutation-induced signaling network rewiring and modulation of signaling dynamics have been systematically characterized (Creixell et al., 2015; Pawson and Warner, 2007), providing a basis for identification of targeted therapies in cancer (Hennessy et al., 2005; Logue and Morrison, 2012). Independently of mutations, kinase overexpression drives tumorigenesis in multiple cancer types and is a critical factor in drug resistance (Eralp et al., 2008; Santarius et al., 2010; Shaffer et al., 2017). Overexpression of phosphatases has been shown to mediate cancer progression and has been linked to the poor prognosis of patients (Julien et al., 2011; Liu et al., 2016; De Vriendt et al., 2013). Overexpression-induced signaling modulation remains largely uncharacterized because factors such as genetic instability induce highly heterogeneous quantities of deregulated signaling proteins in cancer (Abbas et al., 2013), making conventional cell population-based analysis inapplicable. Only recently have technologies emerged that account for such heterogeneity, and can comprehensively quantify signaling network behavior with single-cell resolution. This resolution is required to characterize the abundance-related cellular signaling states and phenotypical alterations caused by a given kinase and phosphatase of interest (Bendall et al., 2011; Budnik et al., 2018; Lun et al., 2017).

Mass cytometry allows simultaneous quantification of over 40 proteins and protein modifications at single-cell resolution and thus enables profiling of complex cellular behaviors in highly heterogeneous samples (Bendall et al., 2011; Bodenmiller et al., 2012; Chevrier et al., 2017). We have recently established and thoroughly validated an approach that couples transient protein overexpression to mass cytometry-based single-cell analysis and have revealed that protein overexpression induces signaling network modulations in an abundance-dependent manner (Lun et al., 2017).

Here, we applied this technique in a human kinome- and phosphatome-wide screen. To achieve this, we generated a library of DNA vectors encoding 541 GFP-tagged kinases and 108 GFP-tagged phosphatases and transfected these vectors into human embryonic kidney HEK293T cells. Single-cell signaling states were determined by simultaneous measurement of 30 phosphorylation sites known to be involved in regulation of growth, proliferation, survival, and stress signaling pathways. Over 10 million individual cells were analyzed in the 659 overexpression conditions with or without 10-minute EGF stimulation, averaging approximately 7,000 measured cells per sample. Assessing the dependence of each measured phosphorylation site on kinase or phosphatase abundance using BP-R^2^, a measure to quantify signaling relationships in single cell analysis (Lun et al., 2017), we identified 1,323 pairs of strong signaling relationships (determined using 108 control experiments). This analysis enabled a functional classification of human kinases and phosphatases based on their abundance-dependent impacts on the signaling network. Furthermore, 208 pairs of previously unknown signaling relationships were identified when compared to the OmniPath database (Türei et al., 2016). By characterizing signaling response dynamics in a follow-up 1-hour EGF stimulation time course, we demonstrated ligand-independent MAPK/ERK activation induced by tyrosine kinase overexpression. We found that in melanoma A375 cells this activation gives rise to BRAF^V600E^ inhibitor resistance. Our screen also revealed that overexpression of ERK-specific phosphatases sustained cell proliferative signals. We confirmed this pro-cancer signaling response using a novel kinase-phosphatase combinatorial overexpression assay.

## Results

### Analysis of the human kinome and phosphatome to study the overexpression effects on signaling network states

Protein abundance variance is often observed in tumors as heterogeneous genomic abnormalities accumulate (Du and Elemento, 2015). We observed up to 50-fold differences in kinase and phosphatase mRNA expression levels among tumor samples in bulk measurements (Figure S1A). This inter-tumoral heterogeneity presumably results in highly variable signaling network states and responses to stimuli or treatments. In addition, a high degree of intra-tumoral expression heterogeneity further challenges cancer therapeutic interventions (McGranahan and Swanton, 2017; Patel et al., 2014). To understand the signaling network modulation in cells that overexpress a defined kinase or phosphatase at various levels, we applied our abundance-dependent signaling network measurement system (Lun et al., 2017) in a kinome- and phosphatome-wide screen.

We cloned open reading frames (ORFs) from the human kinase library (Johannessen et al., 2010) and the human phosphatase library into a vector, enabling expression of GFP-tagged proteins (Couzens et al., 2013). The generated 541 kinase and 108 phosphatase expression clones were transiently transfected into HEK293T cells, individually. Unstimulated cells and cells stimulated for 10 minutes with EGF were harvested and processed with a 126-plex barcoding strategy (adapted from Bodenmiller et al., 2012; Zunder et al., 2015) for simultaneous antibody staining and multiplexed mass cytometry measurement (Figure 1A).

**Figure 1.**
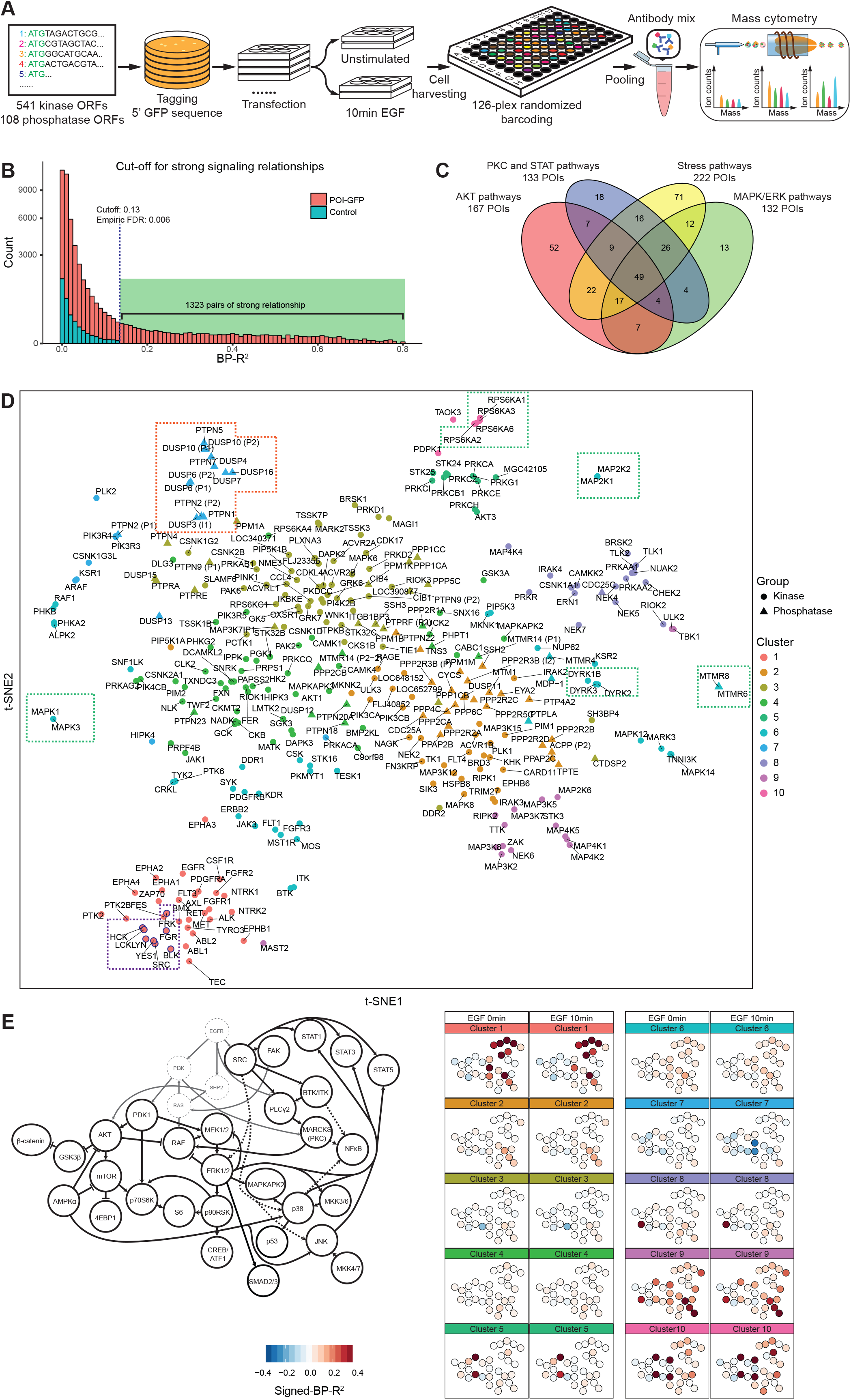
Kinome and phosphatome-wide screen for effects of protein abundance on signaling states and dynamics. **A,** The experimental workflow: ORFs of 541 human kinases and 108 human phosphatases were cloned into a vector to enable expression of GFP-tagged fusion proteins upon transient transfection into HEK293T cells. Cells with or without 10-minute EGF stimulation were harvested, barcoded, and stained with antibody mix before mass cytometry-based single cell analysis. **B,** Plot of counts vs. BP-R^2^ values for control and experimental samples. Cut-off value was determined by analysis of the BP-R^2^ values in all control samples. **C**, Venn diagram showing the quantification of POIs with abundance-dependent influences on the AKT pathway (p-PDK1, p-GSK3β, β-catenin, p-mTOR, p-p70S6K, p-4EBP1, and p-S6), MAPK/ERK pathways (p-RAF, p-MEK1/2, p-ERK1/2, p-p90RSK, p-CREB, and p-SMAD2/3), stress response pathways (p-MKK3/6, p-MKK4/7, p-p38, p-JNK, p-MAPKAPK2, p-AMPKα, and p-p53), PKC pathway and STAT pathways (grouped for illustration purposes; p-SRC, p-FAK, p-BTK, p-PLCγ2, p-MARCKS, p-NFκB, p-STAT1, p-STAT3, and p-STAT5). **D**, t-SNE analysis of overexpressed kinases and phosphatases performed on signed-BP-R^2^ of all measured phosphorylation sites with and without EGF stimulation, color-coded by hierarchical clusters. **E,** The mean signed-BP-R^2^ values of all measured phosphorylation sites in each cluster of kinases or phosphatases shown in a literature-guided canonical signaling network visualization.

Transient transfection generates a gradient of the GFP-tagged protein of interest (POI) expression levels with a range of up to 10^3^-fold enhancement relative to endogenous POI expression range (Lun et al., 2017). The abundance variation of each overexpressed kinase or phosphatase was quantified by mass cytometry with detection by a metal-conjugated anti-GFP antibody. In addition, we simultaneously quantified 30 phosphorylation states of proteins involved in key cancer-related signaling pathways, including the AKT, PKC, STAT, MAPK/ERK, and stress pathways, and five non-signaling markers to indicate cell physiological states (Table S1). Relationship strength between the abundance of GFP-tagged POI and a measured phosphorylation site was quantified by the binned pseudo-R^2^ (BP-R^2^) method (Lun et al., 2017).

We analyzed 108 control samples (FLAG-GFP overexpression or untransfected cells) and used the highest BP-R^2^ score (0.13) of all controls as the cutoff to consider a signaling relationship as “strong”. In total, our human kinome and phosphatome analysis detected 1,323 pairs of strong relationships between POIs and phosphorylation sites (Figure 1B). Among the 649 kinases and phosphatases, 327 had at least one strong signaling relationship (BP-R^2^ > 0.13) to the cancer-related signaling network when overexpressed. Of these, 245 had narrow influences with modulation of one to five signaling nodes, and 26 overexpressed proteins had broad effects on the network with more than ten of measured phosphorylation sites influenced (Figure S1B). We identified 52 kinases or phosphatases that specifically regulated the AKT pathway when overexpressed. Of the 132 POIs that had abundance-dependent effects on the MAPK/ERK pathway, the majority (104 proteins) also initiated cellular stress responses. We also identified 49 proteins that impacted all of the measured signaling pathways, including 11 receptor proteins (e.g., MET, FGFR1, and PDGFRA), and many MAPK cascade activators (e.g., MAP4K1, MAP4K2, and MAP4K5) (Figure 1C).

### Functional classification of kinases and phosphatases based on signaling network modulations

Our kinome and phosphatome screen characterized effects induced by hundreds of POIs on 30 phosphorylation sites in the cancer-related signaling network with and without EGF stimulation (a total of 60 parameters), yielding an unprecedented view on the effects of each kinase and phosphatase. We indicated the sign for signaling relationships (according to their directionality) to the BP-R^2^, generating signed-BP-R^2^ scores that were used for subsequent analyses (Table S2, Methods). To understand the regulatory and functional similarity of overexpressed POIs, we first applied the dimensional reduction algorithm t-SNE (Maaten and Hinton, 2008) to the matrix of all 60 measured signaling parameters (as signed-BP-R^2^ scores) over the 327 signaling network-influential kinases and phosphatases (Figure 1D). As expected, homologous groups (paralogs) of kinases and of phosphatases showed nearly identical influences on signaling and overlapped with each other on the t-SNE plot (Figure 1D, green boxes). This demonstrates that our method sensitively, specifically, and reproducibly detected abundance-modulated signaling behaviors. Interestingly, all the eight overexpressed SRC family members, SRC, YES1, BLK, LCK, LYN, HCK, FGR, and FRK, co-localized in the t-SNE analysis (Figure 1D, purple box), indicating that these kinases have similar functions despite the previously revealed differential patterns of expression (Parsons and Parsons, 2004). Members of protein tyrosine phosphatase (PTPN1, PTPN2, and PTPN5) and dual-specificity phosphatase (DUSP3, DUSP4, DUSP6, DUSP7, DUSP10, and DUSP16) families were grouped together, suggesting similarities in regulating the measured cancer signaling network (Figure 1D, orange box).

We then applied hierarchical clustering based on signed-BP-R^2^ scores of all measured phosphorylation sites (Figure S1C) to analyze functional similarities among all kinases and phosphatases. Based on this analysis we identified 10 major signaling response clusters (color-coded on the t-SNE plot in Figure 1D). Correspondence analysis was performed between these identified clusters and the previously described classes of kinases and phosphatases based on their sequences of the catalytic domain (Johannessen et al., 2010; Sacco et al., 2012a) (Figure S1D). In certain cases, proteins with partial sequence identity had similar influences on signaling. For example, all kinases in cluster 1 were receptor or non-receptor tyrosine kinases (Figure S1D). These kinases were early responders to stimuli, with pleiotropic functions in the signaling networks, as shown in the literature-based graph of canonical EGFR networks (Figure 1E). Clusters 5, 9, and 10 include non-receptor serine/threonine kinases and kinases classified in the group of “other” (i.e., kinases do not fit into any of the major groups) (Figure S1D). Despite conserved catalytic domain sequences, kinases in clusters 5, 9, and 10 induced different cellular responses (Figure 1E). Overexpression of members in cluster 5 activated the PDK1/AKT pathway. Cluster 10 kinases had positive signaling relationships to p90RSK (Ser380), p70S6K (Thr389), PDK1 (Ser241), and MEK1/2 (Ser221) as well as many STAT proteins. Overexpression of cluster 9 components affected proteins involved in the stress response, including MKK4/7 (Ser257/Thr261), p38 (Thr180/Tyr182), and JNK (Thr183/Tyr185). Cluster 9 kinases, when overexpressed, also weakened signaling relationships to MAPK/ERK pathway members, such as MEK1/2 (Ser221), ERK1/2 (Thr202/Tyr204), and p90RSK (Ser380) after EGF stimulation (Figure 1E). Cluster 7 proteins had negative relationships with the signaling mediators of MAPK/ERK pathway when cells were treated with EGF (Figure 1E). Cluster 7 mostly consists of protein tyrosine phosphatases, but also includes a few proteins from the classes of non-receptor serine/threonine kinase and lipid kinases (Figure S1D). Clusters 2, 3, and 4 consist of kinases and phosphatases from multiple sequence-based classes (Figure S1D). This suggests that these proteins induce similar signaling outcomes despite the differences in catalytic domain sequences. In summary, the human kinome- and phosphatome-wide overexpression analysis identified 10 clusters of kinases and phosphatases that partially matched the sequence-based classification. Distinct signaling patterns were found for each cluster.

### Functional enrichment analysis for identified kinase and phosphates clusters

Our analysis indicated that kinases and phosphatases with different catalytic domain sequences could impact signaling networks similarly when overexpressed. To understand this, we performed functional enrichment analysis using the STRING database (Szklarczyk et al., 2017) on the 10 identified clusters (Figure 1D and 2A). We found that seven of the 10 clusters had significant functional enrichment (p < 0.01, Table S3, statistical details in Methods). For each of these clusters, the top five specific functions are shown (Figure 2A). Physical and functional interaction enrichments are shown as protein-protein association networks for clusters 1 and 7 as examples in Figures 2B and 2C and for all other clusters in Figure S2.

**Figure 2.**
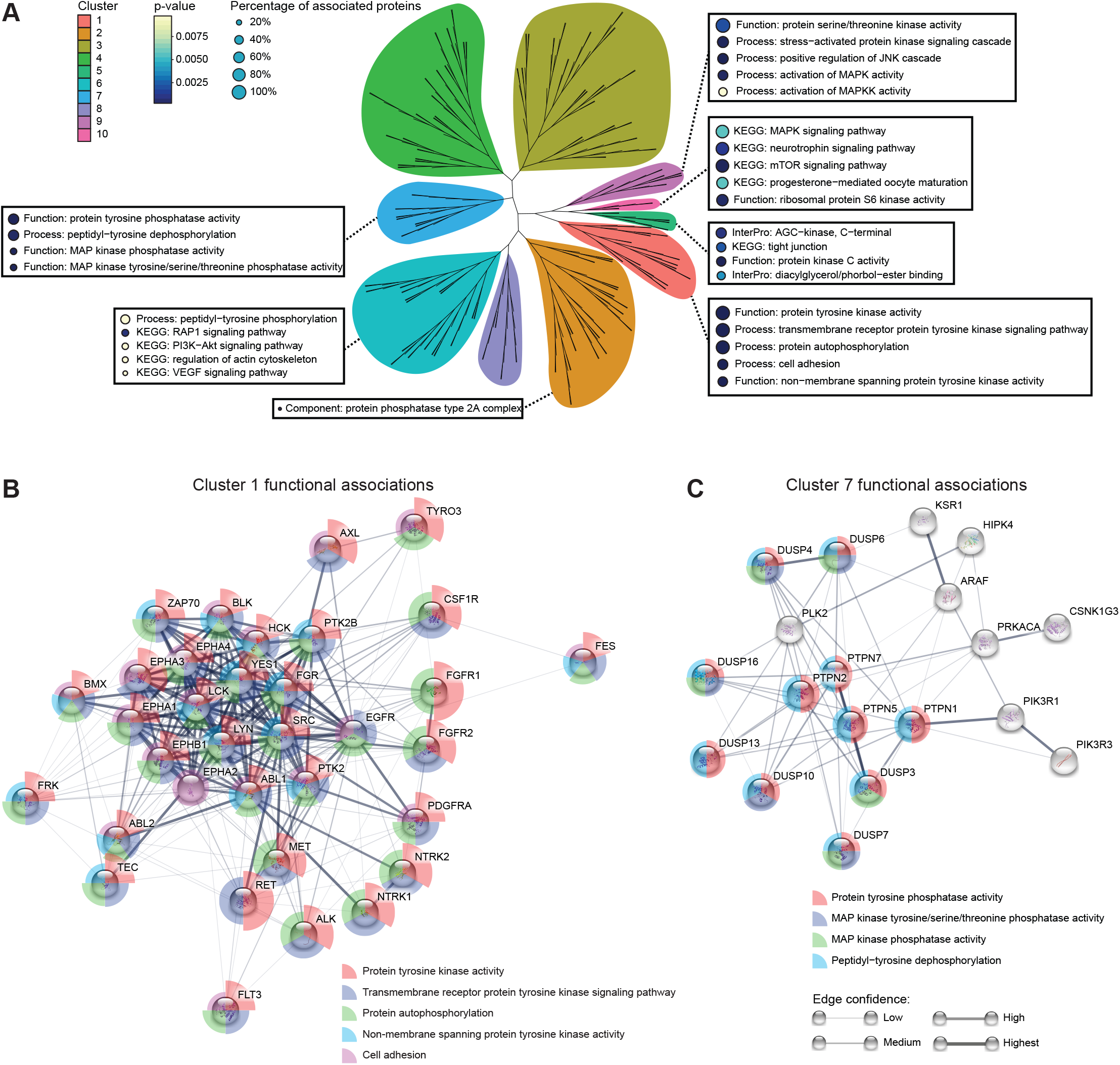
Functional enrichment analysis of kinases and/or phosphatases in each cluster. **A,** Unrooted tree shows the hierarchical clustering of the kinases and phosphatases based on their signed-BP-R^2^ scores. Terms of enriched functions (p<0.05) from each cluster are annotated with circle color indicating the p-value and circle size showing the coverage of cluster components. The percentage of associated proteins is indicated by the size of the adjacent circle. **B-C,** Functional association analysis performed with the STRING database (Szklarczyk et al., 2017) for cluster 1 and cluster 7, respectively. Edges with confidence above 0.2 are shown in the network. Functional enrichments are shown as color-coded-pies with the pie radius indicating the p-value.

As expected, proteins in cluster 1 are enriched for membrane and non-membrane tyrosine kinases with autophosphorylation ability (Figure 2B) with the significance being robust after the removal of redundancy (homology) effects (Table S3, Methods). Although these kinases are closely associated (Figure 2B), we did not find significantly enriched terms for specific signaling pathways or physiological functions (Table S3). Given that all tyrosine kinases of this group generate highly similar signaling events in the measured network (Figure S1C), other factors such as their expression patterns or regulatory protein complexes likely drive their diverse functions in regulating cell phenotypes.

Cluster 7 is enriched for protein tyrosine phosphatases that negatively regulate MAPK pathways. Intriguingly, several kinases are present in this cluster as well, including KSR1 and ARAF, which have similar negative regulatory effects on the MAPK/ERK signaling pathway (Figure 2C). KSR1 and ARAF are core components of the KSR-RAF dimeric protein complex that transduces signal in the MAPK/ERK cascade (Lavoie and Therrien, 2015). Overexpressing one subunit of this protein complex might result in competitive inhibition, diminishing the downstream signal activities in a similar manner as phosphatase overexpression. These analyses demonstrated that proteins with different catalytic functions can mediate highly related signaling responses when overexpressed.

Other clusters were enriched for homologous kinases (clusters 5 and 10) and the protein phosphatase type 2A complex (cluster 2) (Figure S2). We also found that certain kinases associated with the same signaling pathway modulated the measured network differently when overexpressed (Figure S2). Examples are MAPK1 (cluster 6) and RPS6KA1 (cluster 10), which regulate the MAPK/ERK pathway, and PDPK1 (cluster 10) and AKT1 (cluster 4), which regulate the PDK1/AKT pathway. These results suggest that kinase overexpression impacts signaling networks differently to direct kinase activation. Thus, variances in overexpression-induced signaling network modulation are not fully explained by the catalytic functions of kinases and phosphatases. In cells, signaling pathway activity is determined not only by phosphorylation but also by many intrinsic factors, such as protein subcellular localization and protein complex formation.

### Detecting novel signaling relationships from the kinome- and phosphatome-wide analysis

The functions of many kinases and phosphatases analyzed in our screen are unknown or only poorly characterized. We therefore hypothesized that our global analysis could lead to the identification of novel signaling relationships. To assess this, we performed a systematic comparison between all identified overexpression-induced signaling relationships and records in OmniPath, an integrated database of literature-curated signaling interaction information (Türei et al., 2016). We first mapped all pairs of relationships to the OmniPath signaling network, then computed the signed, directed paths for each pair of relationship (Krumsiek et al., 2011; Perfetto et al., 2016). The distance between an overexpressed protein and a measured phosphorylation site is represented by the length of the path (Figure 3A). For example, a distance of zero indicates the relationship between the overexpressed POI and its own phosphorylation levels. Of 14 pairs of signaling relationships with a known distance of zero, with and without 10-minute EGF stimulation, 12 had strong BP-R^2^ values (Figure 3A), revealing that the phosphorylation abundance of kinases is often determined by their own expression level, even in the absence of additional perturbation.

**Figure 3.**
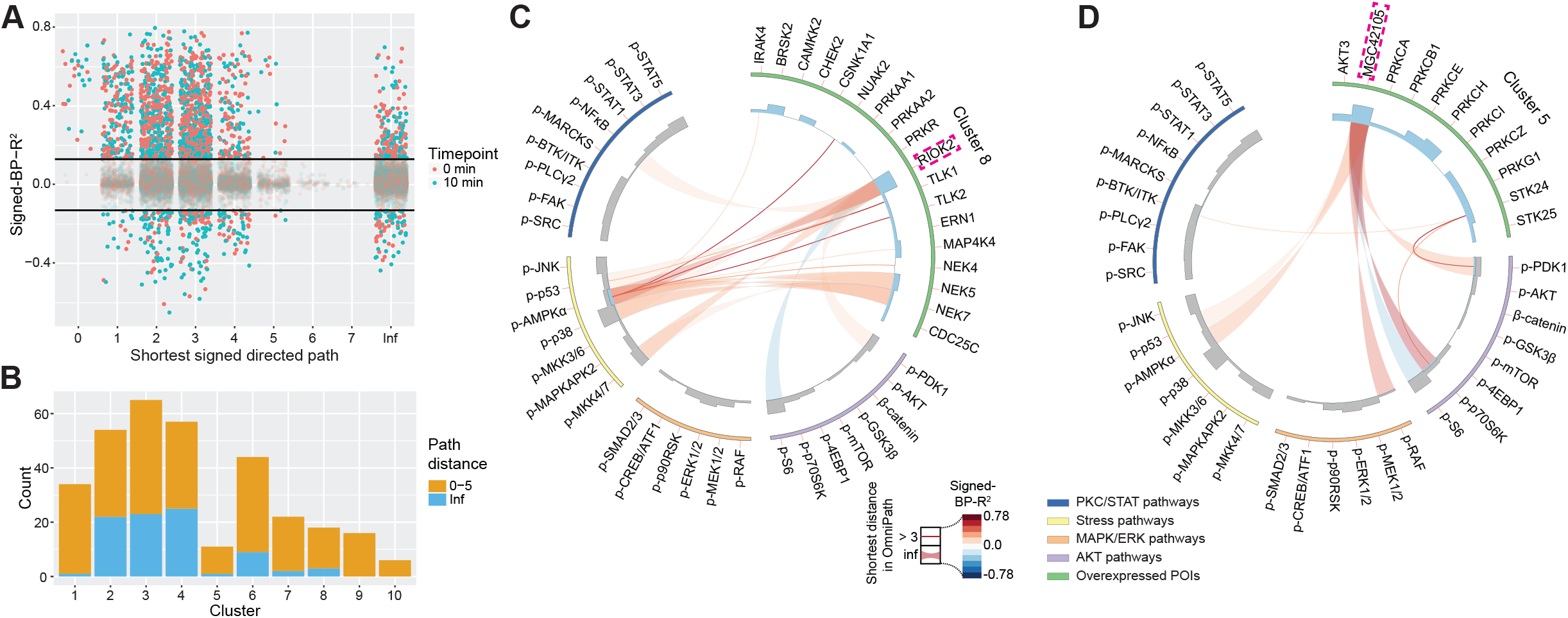
**Prediction of novel signaling connections by comparison with literature evidence in the signaling interaction database OmniPath** (Türei et al., 2016). **A,** Abundance-dependent relationship strength for each pair of overexpressed POI and measured phosphorylation site, as quantified with signed-BP-R^2^, plotted on the length of shortest signed directed path between the two extracted from the OmniPath database. **B,** Occurrences of strong signaling relationships (BP-R^2^ > 0.13) with 0-5 or infinite path length (OmniPath) in each individual hierarchical cluster. **C-D,** For clusters 8 and 5, respectively, shortest signed directed path length for each determined strong signaling relationships shown in Circos plots (Krzywinski et al., 2009).

We detected 208 (16%) strong relationships (BP-R^2^ > 0.13) with infinite distance (Figure 3A, Table S4), indicative of connections not described previously. In total, 93 overexpressed POIs contributed to these signaling relationships, which were enriched (in absolute count) in clusters 2, 3, and 4, and to a less extend in cluster 6 (Figure 3B). We did not detect any relationship with infinite distance in clusters 9 and 10 (Figure 3B); POIs from these clusters participate in MAPK signal transduction (Figure 2A), which is one of the best studied signaling pathways. We also assessed the distribution of infinite paths for each sequence-based kinase and phosphatase class and did not detect any enrichment (Figure S3A). There were also 132 pairs of strong relationships between proteins with length of signed directed path above three in OmniPath, suggesting potentially undiscovered direct or short-range connections (Figure 3A).

Many of the identified novel signaling relationships (i.e., those with infinite path length) and the associated POIs were related to disease and to newly identified kinases (Figures 3C, 3D, S3B, and S3C). For instance, RIOK2 (highlighted in Figure 3C) has been recently shown to correlate with poor prognosis of patients with non-small cell lung cancer, but the underlying signaling mechanisms are unclear (Liu et al., 2016). We discovered that RIOK2 overexpression impacted several phosphorylation sites, most strongly Thr172 on AMPKα, Ser257/Thr261 on MKK4/7, and Thr180/Tyr182 on p38 (Figure 3C), indicating the activation of the AMPK/p38 axis upon RIOK2 overexpression. The AMPK/p38 axis regulates cellular energy metastasis, contributing to cancer cell survival in nutrient-deficient conditions (Chaube et al., 2015; Zadra et al., 2015). We also found that MGC42105 (NIM1K) (highlighted in Figure 3D), a newly identified kinase in cluster 5, regulated the AKT pathway. Overexpression of MGC42105 altered phosphorylation of Ser241 on PDK1, Thr389 on p70S6K, and Ser235/Ser236 on S6. Overexpression of MGC42105 also contributed to the activation of stress pathways, as strong relationships to p-p53 (Ser15) and p-AMPKα (Thr172) were observed (Figure 3D). In summary, mapping our identified signaling relationships to the OmniPath database enabled the identification and assignment of novel signaling functions to many kinases and phosphatases. We also reveal potential novel signaling mechanisms associated with poor prognosis of cancer patients (e.g., for RIOK2).

### In-depth analysis of signaling dynamics reveals overexpression-dependent MAPK/ERK activity

Signaling dynamics are essential for understanding of diseased signaling circuits within a network and in the prediction of drug efficiency (Du and Elemento, 2015; Hughey et al., 2009). We have previously shown that altering expression levels of central signaling proteins in the EGFR signaling network results in complex changes in network dynamics (Lun et al., 2017). Given the key role of signaling dynamics on cell proliferation, growth, and differentiation (Koseska and Bastiaens, 2017), we systematically evaluated kinases and phosphatases from the 10 identified clusters for overexpression-induced signaling dynamic modulations. We calculated the differences in signed-BP-R^2^ scores between the EGF-stimulated (10 min) and unstimulated conditions to identify cases in which overexpression modulates signaling dynamics (i.e., the strength and the shape of abundance-dependent signaling relationship changes between the unstimulated and the 10-minute EGF stimulated conditions). We found that POIs in clusters of 1, 6, 7, 9, and 10 strongly modulated signaling network dynamics when overexpressed (Figure S4A). We then analyzed the overexpression effects of the 39 strongest signaling dynamics influential POIs (criteria described in Methods) over a 1-hour EGF stimulation time course in depth (Figures 4 and S4). The dynamic responses of all measured phosphorylation sites are shown in Figure 4A. Features of the signaling dynamics, such as signaling amplitudes and peak times (see Methods) are shown in Figures S5 and S6.

**Figure 4.**
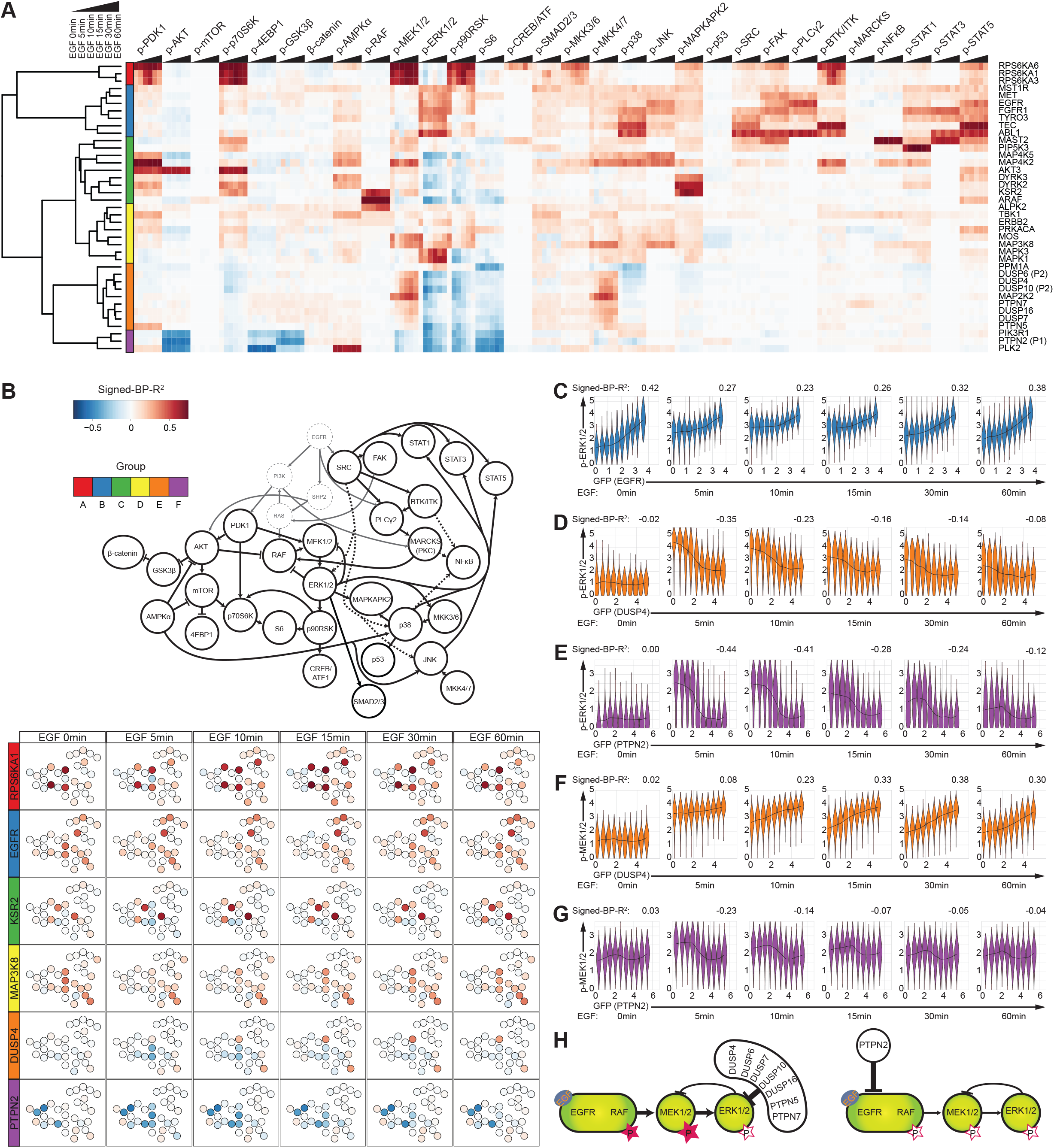
Effects of EGF stimulation on 39 kinases and phosphatases. **A,** Heat map of signed-BP-R^2^ scores for measured signaling relationships over a 1-hour EGF stimulation time course. Six identified groups of kinases and phosphatases are labeled in color codes. **B,** For one representative POI from each group, signaling relationships to all measured phosphorylation sites, as quantified by signed-BP-R^2^, are shown in the literature-guided canonical signaling network map. **C-G,** Violin plots show cell distribution in each of the ten bins divided over the gradients of GFP-tagged POI expression level for EGFR-GFP to p-ERK1/2, DUSP4-GFP to p-ERK1/2, PTPN2-GFP to p-ERK1/2, DUSP4-GFP to p-MEK1/2, and PTPN2-GFP to p-MEK1/2 over the 1-hour EGF stimulation time course. Medians of all 10 bins are connected to indicate the shape of signaling relationships (black lines) with the relationship strength quantified by signed-BP-R^2^, as shown on top of each individual plot. **H,** Schematic illustration of how two sets of phosphatases induce different abundance-dependent influences on the signaling dynamics of the MAPK/ERK cascade.

Hierarchical clustering of the overexpression-induced EGFR signaling dynamics of the 39 selected proteins revealed six groups (Figure 4A). Each of these six groups typically reflected one or two of the 10 major signaling response clusters identified in the previous sections (Figure 1D); the correspondence is shown in Figure S4B. Signaling network responses for one representative kinase or phosphatase from each of the six identified groups are illustrated in Figure 4B. Interestingly, the EGF stimulation-influenced relationships were mostly observed for phosphorylation sites within the MAPK/ERK signaling cascade, including p-MEK1/2 (Ser221), p-ERK1/2 (Thr202/Tyr204), p-p90RSK (Ser380), and p-S6 (Ser235/Ser236), rather than AKT, PKC, STAT, or stress response pathways (Figure 4A). Given that the MAPK/ERK is the major proliferative pathway that is known to be involved in tumor progression and drug response, we focused our subsequent analyses on this pathway to identify novel regulatory mechanisms potentially relevant to cancer.

EGFR overexpression (as an example of group B) mediated activation of the MAPK/ERK pathway (Figure 4B), multiple STAT kinases, PLC/2, and the stress signaling pathways. Phosphorylation of Thr202/Tyr204 on ERK1/2 was elevated in cells with high levels of GFP-tagged EGFR in the absence of EGF stimulation (Figure 4C). These cells responded weakly to EGF stimulation, indicating ligand-independent ERK activation (Figures 4C and S5B). Overexpression of other members of group B, including MST1R, MET, FGFR1, TYRO3, TEC, and ABL1, influenced signaling dynamics of p-ERK1/2 and p-90RSK in a manner similar to that of EGFR overexpression (Figure 4A). These proteins correspond to the global cluster 1 (Figure S4B), and members of this class mediate oncogenic signaling for many cancer types (Duan et al., 2016; Paul and Mukhopadhyay, 2004; Salgia, 2017). Taken together, this suggests that ligand-independent MAPK/ERK signaling activation causes uncontrolled cell proliferation.

Interestingly, in the absence of EGF stimulation, overexpression of phosphatases DUSP4 (from group E) and PTPN2 (from group F) did not affect levels of phosphorylation in the MAPK/ERK pathway (Ser221 on MEK1/2, Thr202/Tyr204 on ERK1/2, or Ser380 on p90RSK) (Figures 4B, 4D-4G, S4C, and S4D). This suggests either a mechanism compensates for phosphatase overexpression to maintain basal MAPK/ERK signaling or the overexpressed phosphatases are inactive without EGF stimulation. Upon EGF stimulation, signaling dynamics on phosphorylation sites of the MAPK/ERK pathway were modulated by DUSP4 and PTPN2 in an abundance-dependent manner, as negative signaling relationships to p-ERK1/2 and p-p90RSK were detected (Figures 4B, 4D, 4E, S4C and S4D). DUSP4 or PTPN2 overexpression also resulted in reduced signaling amplitudes on p-ERK1/2 and p-p90RSK (Figure S5C). Intriguingly, different p-MEK1/2 dynamics were observed in cells overexpressing these two phosphatases. There was a strong positive relationship between GFP-tagged DUSP4 and p-MEK1/2 beginning at 10 minutes after EGF addition (signed-BP-R^2^ = 0.23) with relationship strength constantly increasing until the 30-minute time point (Figure 4F), indicating a more sustained MEK1/2 phosphorylation in cells with higher levels of DUSP4 than in cells with lower DUSP4 expression. In contrast, a negative relationship between GFP-tagged PTPN2 and p-MEK1/2 was observed during EGF stimulation with a strong relationship at the 5-minute (signed-BP-R^2^ = −0.23) and 10-minute (signed-BP-R^2^ = −0.14) time points (Figure 4G). DUSP4 is known to selectively de-phosphorylate ERK1 and ERK2 (Guan and Butch, 1995). Our data indicate that during EGF stimulation, DUSP4 overexpression diminishes ERK1/2 phosphorylation, and, in turn, the negative feedback from ERK1/2 to MEK1/2 is likely attenuated, resulting in constant activation of MEK1/2. Substrates of PTPN2 are primarily membrane kinases including EGFR (Mattila et al., 2005). Overexpression of PTPN2, therefore, downregulates activation states of all measured signaling proteins known to be downstream of EGFR, including MEK1/2 and ERK1/2 (Figure 4h). Systematic analysis of all overexpressed phosphatases over the 1-hour time course after EGF addition confirmed that all other phosphatases in group E (DUSP6, DUSP7, DUSP10, DUSP16, PTPN5, and PTPN7) target phosphorylation sites of Thr202/Tyr204 on ERK1/2, thereby decreasing the negative feedback from ERK1/2 to MEK1/2 and causing sustained MEK1/2 activation (Figures 4A and 4H).

In conclusion, these signaling dynamic analyses indicate that first, overexpression of tyrosine kinases including EGFR induce EGF ligand-independent MAPK/ERK signaling activation. Second, phosphatases differentially regulate MAPK/ERK signaling responses determined by their substrate specificities. Third, the ERK-specific phosphatases control the strength of the negative feedback loop from ERK1/2 to MEK1/2 in an abundance-dependent manner.

### Pairwise overexpression analysis reveals phosphatase sustains the kinase-induced MAPK/ERK signaling

Phosphatase overexpression is oncogenic in different tumor types, but the signaling mechanisms remain unclear (Julien et al., 2007, 2011). Recent work indicates that overexpressed phosphatases increase the malignancy of cancers that have a hyper-activated MAPK/ERK pathway (Julien et al., 2007; Low and Zhang, 2016; De Vriendt et al., 2013). Our result in the previous section suggested a mechanism through which overexpression of ERK-specific phosphatases sustains MEK phosphorylation levels (Figures 4F and 4H). To assess whether an additional, secondary signaling input that increases MAPK pathway activity could lead to oncogenic-like signaling, we developed a combinatorial transfection assay in which overexpression of a kinase and a phosphatase were detected via a FLAG-tag and a GFP-tag, respectively, providing a two-dimensional analysis of abundance-dependent signaling modulation on the single cell level (Figure 5A). Using this approach, we analyzed the MAP2K2, MAPK1, and RPS6KA1 (also known as MEK2, ERK2, and p90RSK1) kinases, and the DUSP4, DUSP7, and PTPN2 phosphatases in nine combinations of double overexpression over a 1-hour EGF stimulation time course (Figure 5B). After measurement, we subdivided cells according to the expression levels of the FLAG-tagged kinase and GFP-tagged phosphatase into 25 bins within the two-dimensional overexpression space. For each bin, signaling states as defined by phosphorylation levels, and signaling trajectories with respect to all individual bins over the 1-hour EGF time course were analyzed.

**Figure 5.**
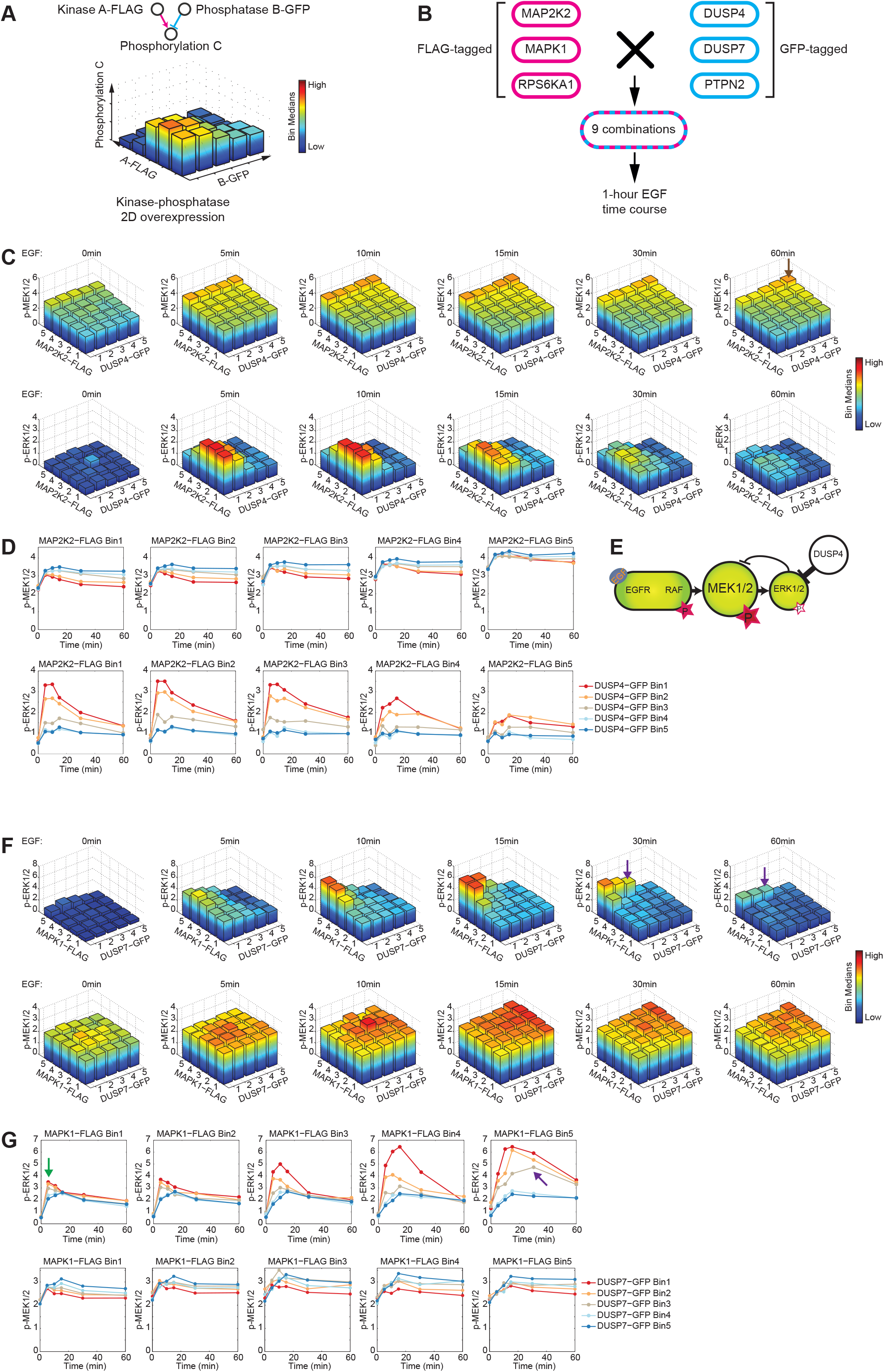
Pairwise overexpression of two-dimensional protein abundance dependency analysis of signaling behaviors. **A,** Workflow of the pairwise overexpression. Two plasmids encoding a FLAG-tagged kinase and a GFP-tagged phosphatase were transfected into HEK293T cells successively. Cells were binned into 25 groups according to their FLAG and GFP abundances. The median level of each measured phosphorylation site was computed for each bin. **B**, Kinases MAP2K2, MAPK1, and RPS6KA1 and phosphatases DUSP4, DUSP7, and PTPN2 were selected for the pairwise overexpression, generating nine overexpression combinations in total. **C,** In cells with pairwise overexpression of a FLAG-tagged kinase MAP2K2 (MEK2) and a GFP-tagged phosphatase DUSP4, median phosphorylation levels of p-MEK1/2 and p-ERK1/2 are plotted for all bins over the 1-hour EGF stimulation time course. **D,** Signaling trajectories of p-MEK1/2 and p-ERK1/2 plotted as the medians of each individual bin over the 1-hour EGF stimulation time course. **E,** Schematic illustrating the modulation of RAF/MEK/ERK cascade signaling states and dynamics by the pairwise overexpression. **F-G,** Analysis of dependence of p-ERK1/2 and p-MEK1/2 signaling on MAPK1(ERK2)-FLAG and DUSP7-GFP.

When overexpressed individually, we observed that DUSP4 overexpression sustained the phosphorylation of Ser221 on MEK1/2 over the 1-hour EGF stimulation time course likely due to the weakened ERK to MEK negative feedback (Figures 5C and 5D). MAP2K2 (MEK2)-FLAG overexpression led to an increased MEK1/2 phosphorylation (Figure 5C). Interestingly, combined signaling inputs from MAP2K2-FLAG and DUSP4-GFP co-overexpression further increased the hyper-activated states of MEK1/2 over the 1-hour EGF stimulation time course compared to the activation induced by MAP2K2-FLAG overexpression alone (Figures 5C-5E, brown arrow). Moreover, in cells with simultaneously overexpressed MAP2K2-FLAG and DUSP4-GFP, the downstream ERK1/2 phosphorylation on Thr202 and Tyr204 were inhibited (Figures 5C-5E). Highly activated MEK1/2 could lead to ERK-independent oncogenic-like signaling as revealed previously (Burgermeister and Seger, 2008; Takahashi-Yanaga et al., 2004). In contrast, double overexpression of MAP2K2-FLAG and PTPN2-GFP (the latter a phosphatase targeting EGFR) did not sustain the MEK1/2 signaling; rather, it dampened the MEK1/2 signaling amplitudes in response to EGF stimulation (Figure S7).

Overexpression of FLAG-tagged MAPK1 (ERK2) drastically augmented ERK1/2 phosphorylation state during EGF stimulation (Figure 5F), increased p-ERK1/2 amplitudes, and delayed p-ERK1/2 peak times (Figure 5G). These results are in agreement with findings of previous studied MAPK1 overexpression effects (Lun et al., 2017). The simultaneous overexpression of MAPK1-FLAG and DUSP7-GFP decreased p-ERK1/2 levels at all time points and reduced the signaling amplitudes. Further, DUSP7 delayed p-ERK1/2 peak times upon EGF stimulation: in cells with the highest MAPK1 abundance and mid-level overexpression of DUSP7, ERK1/2 phosphorylation peaked at 30 minutes after EGF addition (Figures 5F and 5G, purple arrow), whereas in untransfected cells, p-ERK1/2 peaked at the 5-minute time point (Figures 5F and 5G). As expected, DUSP7 overexpression also resulted in constant MEK1/2 phosphorylation (Figures 5F and 5G, green arrow). Compared to cells only overexpressing MAPK1 (ERK2), which induced strong but transient ERK activation, the additional low-to-mid levels of DUSP7 decreased the ERK1/2 phosphorylation amplitude and partially limited the negative feedback signal from ERK to MEK, inducing a sustained MEK activation and prolonged ERK signal.

Assessing signaling responses in the pairwise overexpression assay, we demonstrated kinase and phosphatase coregulatory mechanisms in the MAPK/ERK cascade. We confirmed that phosphatase overexpression does not dampen signaling activity at the steady states, even with activated oncogenic signaling. Further, our analysis indicates that the overexpression of certain phosphatases, such as DUSP4 and DUSP7, lead to sustained activation of ERK due to the reduced negative feedback strength. This mechanism might underlie the pro-cancer effects of phosphatase overexpression.

### Signaling relationship to p-ERK1/2 predicts overexpression-induced vemurafenib resistance in melanoma A375 cells

As protein overexpression has been correlated with drug resistance of cancer cells (Johannessen et al., 2010; Shaffer et al., 2017), we next sought to determine whether our kinome- and phosphatome-wide signaling network profiles could identify kinases or phosphatases that, when overexpressed, induce drug resistance; these enzymes can be potential biomarkers of drug resistance. In melanoma cells carrying the BRAF^V600E^ mutation, overexpression of certain kinases is associated with *de novo* or acquired resistance to RAF inhibition; Johannessen et al. has identified nine kinases that drive resistance when overexpressed in a cell-based assay (Johannessen et al., 2010). Eight of these proteins were analyzed in our screen, and interestingly, six had abundance-dependent signaling modulations to p-ERK1/2 (Thr202/Tyr204) in unstimulated cells (Figure S8A). The 10-minute EGF stimulation reduced relationship strengths for each of these six kinases (Figure S8B), indicating that these overexpression-related ERK activations were ligand-binding independent (i.e., cells with high POI-GFP levels did not respond to EGF stimulation, Figure 4C). These data suggest that overexpression of kinases that induce ligand-independent MAPK/ERK pathway activation might underlie resistance to BRAF^V600E^-targeted inhibitors.

To determine whether signaling relationships to p-ERK1/2 in our kinome- and phosphatome-wide analysis have potential as biomarkers for drug resistance, we examined kinases in our screen with the highest signed-BP-R^2^ values in relation to p-ERK1/2 in the absence of EGF stimulation, including ABL1, BLK, FES, MAP3K2, MAP3K8, MOS, NTRK2, SRC, and YES1. These kinases, and MEK1^DD^ as positive control, were overexpressed in A375 cells that express BRAF^V600E^. Cells were subsequently treated for 48 hours with the BRAF^V600E^ inhibitor vemurafenib or DMSO as a control (Figure 6A, all experiments performed in three replicates). In A375 cells, the BRAF substrate MEK1/2 is activated and basal levels of p-ERK1/2 are high. As expected, overexpression of MEK1^DD^, which constitutively phosphorylates Thr202 and Tyr204 on ERK1/2 (Johannessen et al., 2010; Lun et al., 2017), only slightly enhanced ERK1/2 phosphorylation in cells treated with DMSO (Figure 6B). A weak signaling relationship was observed between GFP-tagged MEK1^DD^ and p-ERK1/2 with a signed-BP-R^2^ of 0.11 (Figure 6B, left). Compared to the DMSO control, treatment with vemurafenib inhibited BRAF^V600E^ activity and reduced p-ERK1/2 levels in untransfected cells and in cells with low MEK1^DD^ expression levels. However, cells with high levels of MEK1^DD^ were insensitive to vemurafenib, such that hyper-phosphorylated ERK1/2 remained after the 48-hour treatment (signed-BP-R^2^ value of 0.53, Figure 6B, right). In the control cells without MEK^DD^ overexpression, the effect of vemurafenib on p-ERK1/2 levels did not alter signaling relationships as quantified by signed-BP-R^2^ (Figures S8C and S8D).

**Figure 6.**
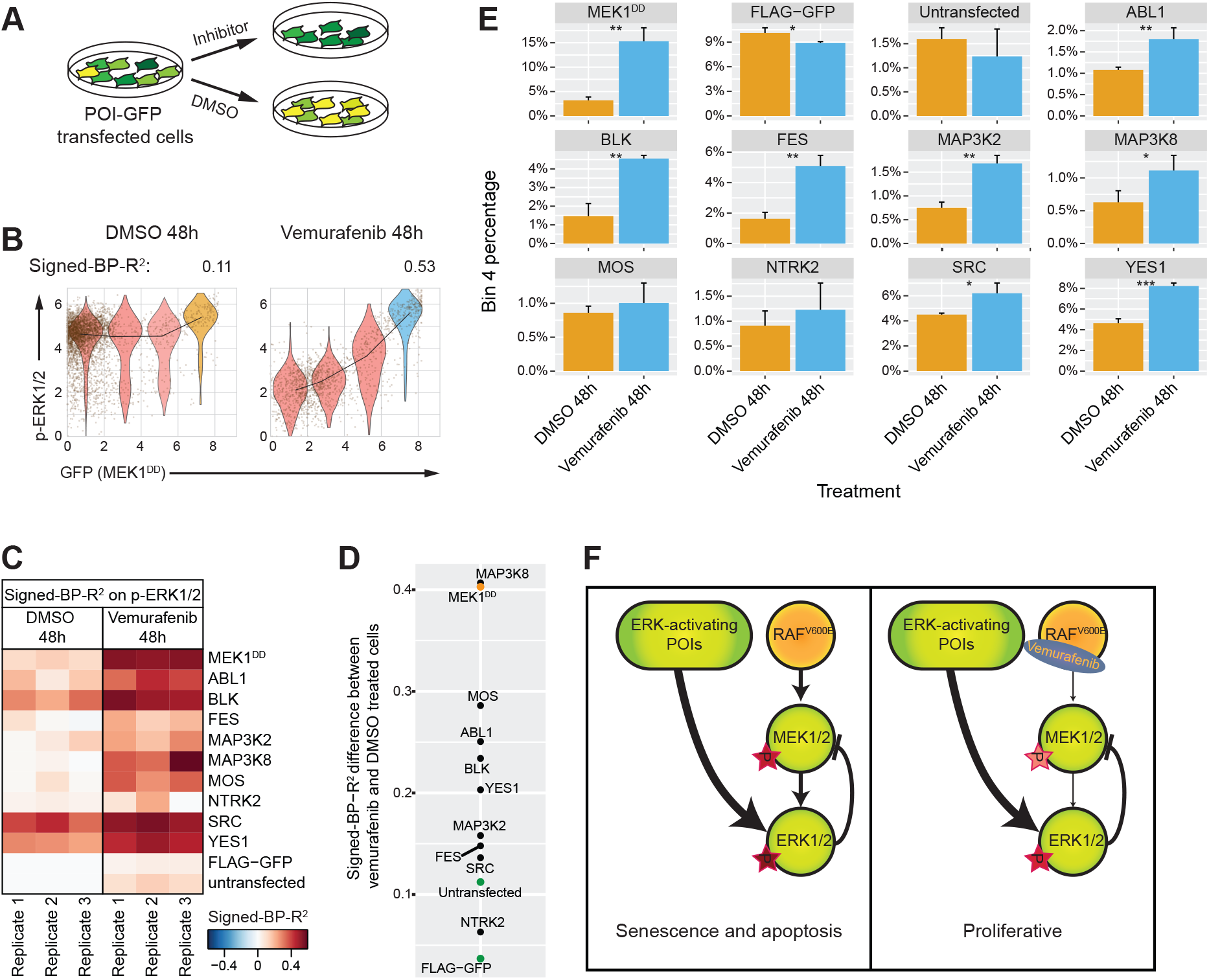
Kinase overexpression induces vemurafenib resistance in the melanoma A375 cells by activating ligand-binding independently of ERK signaling. **A,** Selected kinases were transfected into A375 cells and cultured in the inhibitor- or DMSO-containing media for 48 hours before the signaling states and enrichment of kinase overexpression cells were assessed. **B,** Data from each sample were divided into four bins depending on the expression level of the GFP-tagged kinase. Signed-BP-R^2^ analysis was performed to quantify signaling relationships with and without vemurafenib. **C,** Histogram of abundance-dependent relationship strength for each overexpressed POI to p-ERK1/2, quantified as signed-BP-R^2^ from three replicate experiments. **D,** The mean differences of the three replicates between vemurafenib-treated cells and DMSO-treated cells in their signed-BP-R^2^ scores with p-ERK1/2. **E,** Proportion of cells in bin 4, cells with the highest levels of POI, for each individual overexpressed kinase in vemurafenib-treated cells and DMSO-treated cells. (* p < 0.05; ** p < 0.01; *** p < 0.001, n = 3) **F,** Illustrations of how ERK-activating POIs maintain the proliferative state in cells treated with vemurafenib and, in the un-drugged condition, how (despite overexpression of these POIs) over-activated ERK signaling leads to cell senescence and apoptosis.

Similar as observed in cells overexpressing GFP-tagged MEK1^DD^, vemurafenib treatment also increased signaling relationships between all our analyzed GFP-tagged kinases (ABL1, BLK, FES, MAP3K2, MAP3K8, MOS, NTRK2, SRC, and YES1) and p-ERK1/2 (Figure 6C). A closer inspection of the signed-BP-R^2^ signaling strength of cells treated for 48 hours with vemurafenib and DMSO-treated cells showed that eight of the nine tested kinases had higher values than those of negative controls (Figure 6D). These results indicate that overexpression of ERK-activating kinases limits the effects of vemurafenib in reducing p-ERK1/2 signaling, similarly as the MEK^DD^ positive control.

To assess the survival of cells with different expression levels of each kinase, we assigned each single cell into one of four bins based on the abundance of GFP-tagged POI and calculated the percentage of the number of cells in each bin relative to the total cell count (Figure 6B). As expected, the positive control cells that expressed MEK1^DD^-GFP had significant cell enrichment in the fourth bin (i.e., the bin with the highest expression level of the GFP-tagged POI), with four times higher percentage of cells at this level after 48 hours of vemurafenib treatment compared to the DMSO-treated control (Figure 6E). Similarly, all nine examined kinases showed enrichment of cell abundance in the fourth bin: in six cases, this enrichment was statistically significant (Figure 6E).

These results indicate that, in melanoma A375 cells, overexpression of kinases capable of ligand-independent ERK activation reduces cellular dependency on signaling inputs from BRAF^V600E^ for proliferation, rendering the cells less sensitive to the BRAF^V600E^ inhibitor vemurafenib (Figure 6F). In the absence of inhibitor, however, cells overexpressing ERK-activating POIs do not acquire proliferative advantage (Figure 6F). As has been shown previously, the hyperactivity of ERK induces cell senescence and apoptosis (Cagnol and Chambard, 2010; Xu et al., 2014). Our kinome and phosphatome screen detected 54 POIs that, when overexpressed, caused EGF-independent ERK activation (Table S5). These proteins can potentially be used as biomarkers to predict resistance to BRAF^V600E^ inhibitors in melanoma patients carrying BRAF mutations.

## Discussion

The data described here are unique for the broad coverage of the human kinome and phosphatome, the multiplexed measurement of cellular phosphorylation states and dynamics at single-cell resolution, and the wide abundance range (over three orders of magnitude) over which proteins of interest were studied. Our analyses enabled protein abundance-determined functional classification, signaling kinetics quantification, and the identification of potential biomarkers of drug resistance.

Extending our previously established approach (Lun et al., 2017) for analysis of the dependence of signaling behaviors on protein expression levels to the human kinome and phosphatome, we transiently transfected a library of 649 plasmids encoding GFP-tagged kinases and phosphatases individually into HEK293T cells, yielding gradient overexpression levels for each GFP-tagged POI. The abundance of each overexpressed kinase or phosphatase was quantified simultaneously with 30 cancer-related phosphorylation sites and five non-signaling markers (Table S1) using a mass cytometry-based multiplexed single-cell assay. Signaling states and dynamics over the expression continuum for every analyzed POI were comprehensively profiled. We applied the recently developed statistical measure, BP-R^2^, to quantify signaling strength between each GFP-tagged POI and each measured phosphorylation site (Lun et al., 2017) and classified all overexpressed kinases and phosphatases based on their abundance-determined signaling network states with or without 10-minute EGF stimulation.

Protein abundance and mRNA expression levels of kinases and phosphatases have been quantified in normal and diseased tissues by multiple approaches (Petryszak et al., 2016; Uhlen et al., 2017; Wang et al., 2015). Our analysis, for the first time, characterized at kinome- and phosphatome-wide scope how these proteins differentially modulate network behaviors when expressed over a concentration gradient. In the overexpression effect-based functional classification, we assigned kinases and phosphatases into 10 clusters, each with a distinct function in signaling transduction. These clusters only partly agreed with the kinase and phosphatase classes based on their sequence of catalytic domain, indicating the dissimilar network alterations between signaling protein overexpression and activation. Sequence-based classification considers the catalytic specificities of kinases and phosphatases. However, by altering the signaling protein expression levels (or concentration), dynamics of both upstream and downstream signaling events, and the assembly of multiprotein complexes can be modulated, resulting in more complicated network changes.

Interestingly, our kinase and phosphatase classification and the functional enrichment analysis suggest that kinases, such as KRS1 (a pseudo-kinase) and ARAF, negatively regulate MAPK/ERK signaling when overexpressed, similarly to many tyrosine phosphatases (Figures 1D, 2A, and 2C). In the MAPK/ERK cascade, dimerization between KSR proteins (KSR1 and KSR2) and RAF proteins (ARAF, BRAF, and CRAF), is required for the activation of the downstream kinase MEK (Lavoie and Therrien, 2015; Rajakulendran et al., 2009). Overexpressing either of KSR1 or ARAF leads to competitive protein binding that disrupts the formation of the dimeric protein complex. Interestingly, cells with CRAF (also known as RAF1) or KSR2 (both in cluster 6) overexpression showed less degree of MAPK/ERK signaling attenuation (Figure S1C), compared to those with KSR1 or ARAF overexpression. This suggests the differential capability of individual RAF proteins or KSR proteins in forming homodimers that can compensate overexpression-induced signaling disruption.

Phosphatase overexpression is observed to drive tumor progression with unclear signaling mechanisms (Julien et al., 2011; Low and Zhang, 2016; De Vriendt et al., 2013). We indicated that rather than directly activating a cancer-driven signaling pathway, overexpression of ERK-specific phosphatases modulates signaling dynamics that leads to prolonged proliferative signal in cells. Phosphatases have been suggested in many recent studies as potential therapeutic targets (Bollu et al., 2017; Julien et al., 2011; Low and Zhang, 2016). Our result indicates again the importance of developing phosphatase inhibitors for cancer treatment.

Following a pilot kinome study on overexpression-related resistance to RAF inhibition (Johannessen et al., 2010), we discovered that ligand-independent ERK activation induced by kinase overexpression to be the underlying signaling mechanism leading to above drug resistance. Our kinome and phosphatome analysis further identified 54 proteins that caused ligand-independent ERK activation when overexpressed. These proteins were then suggested as potential biomarkers for RAF inhibitor resistance. Mass cytometry-based single cell analysis enables identifying differed signaling responses to a drug treatment in rare cell populations. It is therefore more sensitive in identifying biomarkers for overexpression-induced drug resistance, compared to the previous population-based assay (Johannessen et al., 2010). Our data can be further used to reveal signaling mechanisms in diseased cellular behaviors that are caused by kinase or phosphatase overexpression. Then, following the discovered pathological signaling behaviors, more biomarkers can be identified for the given disease, based on our kinome- and phosphatome-wide analysis.

Our analysis has several limitations. First, the measured overexpression effects can be indirect (i.e., a protein overexpression might lead to cellular stress that activates MAPK/p38 and MAPK/JNK cascades). However, indirect effects are also important signaling responses that can be indicative for any cells with such overexpression. Second, our mass cytometry-based analysis applies pre-selected antibodies targeting 30 specific phosphorylation sites that does not capture all signaling responses induced by overexpression of a kinase or a phosphatase. Nevertheless, our antibody panel is designed based on literature information to cover the most critical and informative phosphorylation sites in the cancer-related signaling network. Third, the GFP-tag can potentially disrupt the function of a kinase or phosphatase. Likely, a non-functional protein does not induce specific signaling network modulation and should therefore yield weak BP-R^2^ values (≤ 0.13) only.

In summary, we demonstrated, in the human kinome- and phosphatome-scale analysis, how overexpression of each signaling protein modulates signaling networks in an abundance-dependent manner. This establishes that protein expression levels can result in different signaling states in a heterogeneous population. Our analysis expands the functional classification of the human kinases and phosphatases, and suggests 208 novel signaling relationships that can be interrogated to improve our understanding of signaling causality and network structure. We showed novel oncogenic-like signaling mechanisms and identified cancer biomarkers with our analysis. Our data are also suitable for the inference of signaling pathway kinetics using mathematical models and for the development of network reconstruction methods.

## Author contributions

X.-K.L. and B.B. conceived the study. X.-K.L. performed experiments, data processing, and data analysis. D.S. performed functional enrichment and functional association analysis. A.G. performed novel relationship analysis. N.D. cloned the phosphatase expression library. V.Z. implemented the BP-R^2^ platform and helped with data analysis. X.-K.L., D.S., A.G., N.D., V.Z., J.S.-R., C.M., and B.B. performed the biological interpretation. X.-K.L. and B.B. wrote the manuscript with input from all authors.

## Acknowledgement

We would like to thank the Bodenmiller lab for support and fruitful discussions, the Sommer laboratory for sharing experimental materials, the Lehner lab and the Mosimann lab for sharing equipment, Dr. Vinko Tosevski and Dr. Tess Brodie at the Mass Cytometry Facility, University of Zürich for support and troubleshooting help. We would especially like to thank Dr. A.-C. Gingras, Lunenfeld-Tanenbaum Research Institute, for sharing the pDEST vectors used in this study. This work was supported by an SNSF R`Equip grant, a SNSF Assistant Professorship grant (PP00P3-144874), by the European Research Council (ERC) under the European Union’s Seventh Framework Program (FP/2007-2013)/ERC Grant Agreement n. 336921 and an NIH grant (UC4 DK108132).

